## Supplementary Figure for "Global Endometrial DNA Multi-omics Analysis Reveals Insights into mQTL Regulation and Associated Endometriosis Disease Risk"

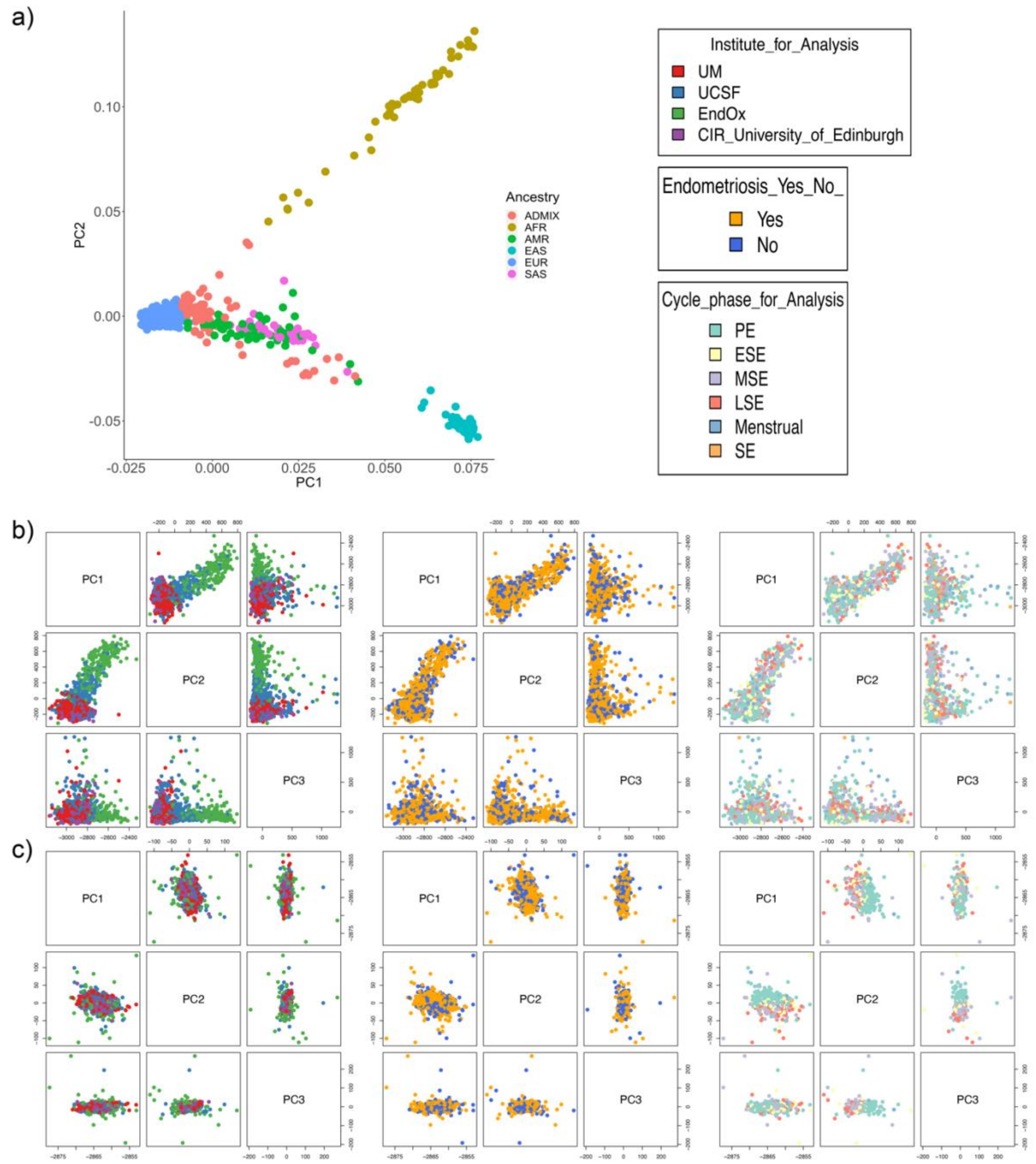


**Supplementary Figure 1. PCAs of DNA Methylation and Genotyping.** a) Genotype data PCA plots for PC1 vs. PC2 with genetic ancestry of samples shown by the point colours. b) PCA plots of methylation pre-batch correction coloured according to institute (left), case:control (middle) and cycle phase (right). c) PCA plots of methylation post-batch correction coloured according to institute (left), case:control (middle) and cycle phase (right).

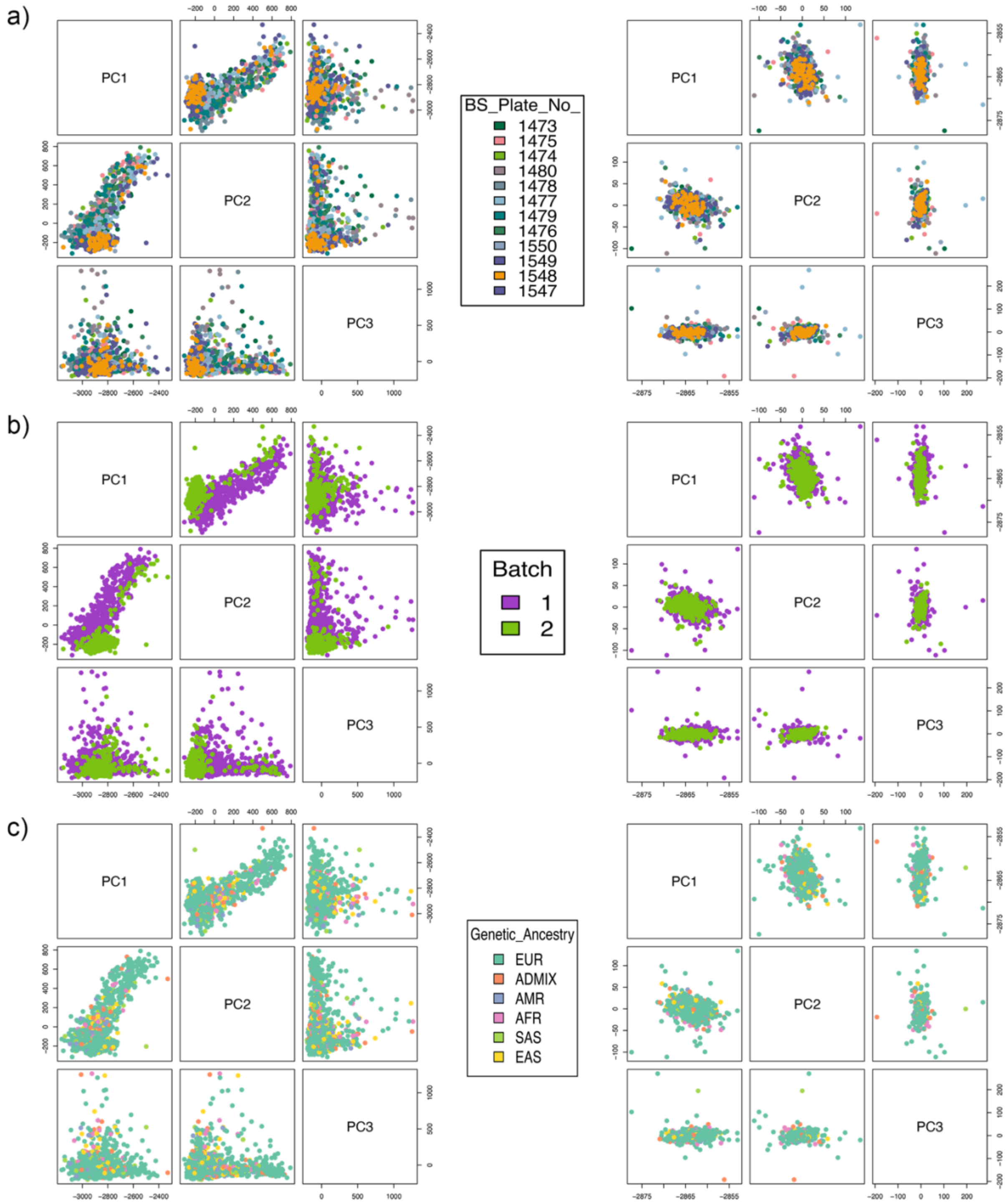


**Supplementary Figure 2. Methylation Data PCs Before and After Batch Correction.** a) PCA plots of methylation coloured according to plate pre-batch correction (left) and post-batch correction (right). b) PCA plots of methylation coloured according to processing batch pre-batch correction (left) and post-batch correction (right). c) PCA plots of methylation coloured according to genetic ancestry pre-batch correction (left) and post-batch correction (right).

a)

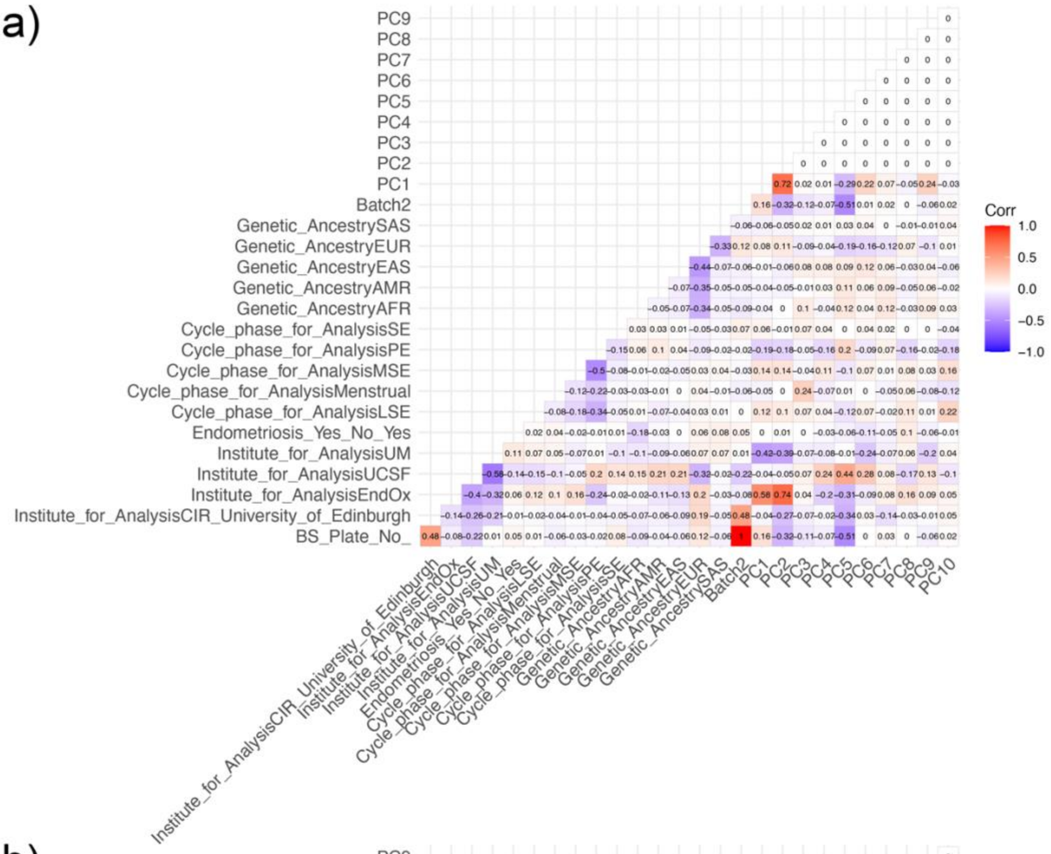


b)

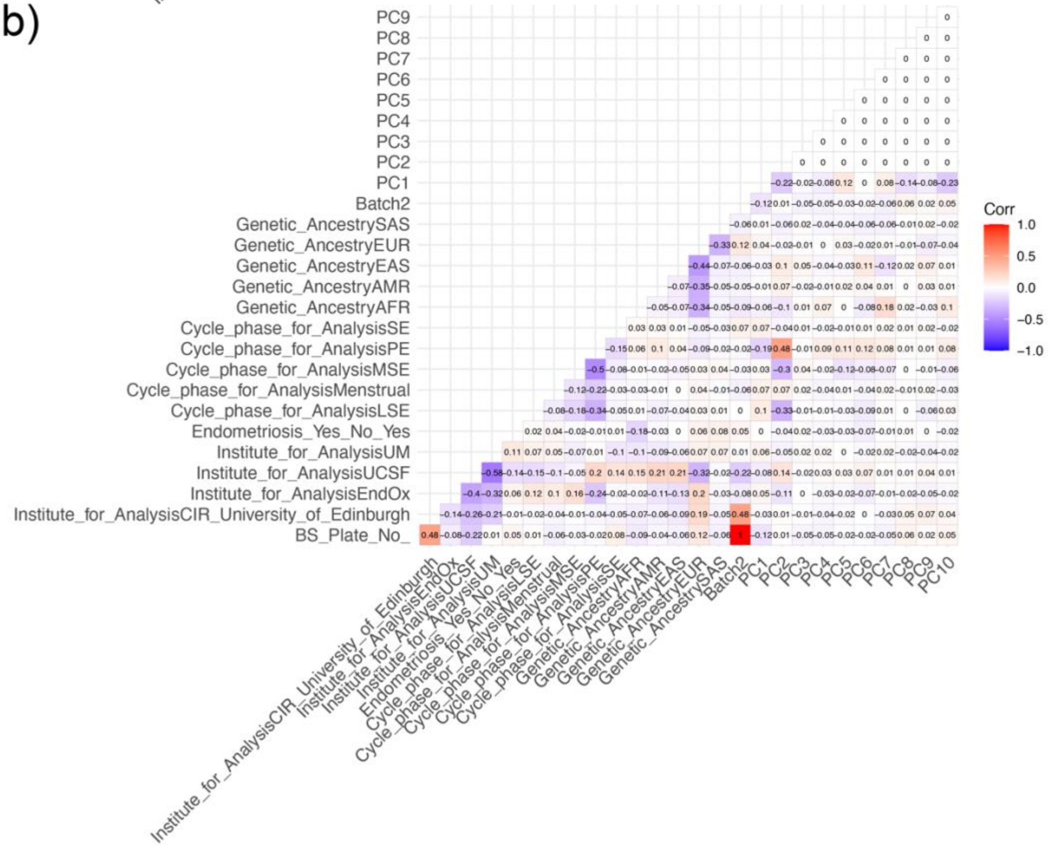


**Supplementary Figure 3. Covariates and Principle Component (PC) Correlation Plots.** a) Correlation between the top PCs and sample variables before correction with surrogate variables (SVs). b) Correlation between the top PCs and sample variables after correction with surrogate variables (SVs).

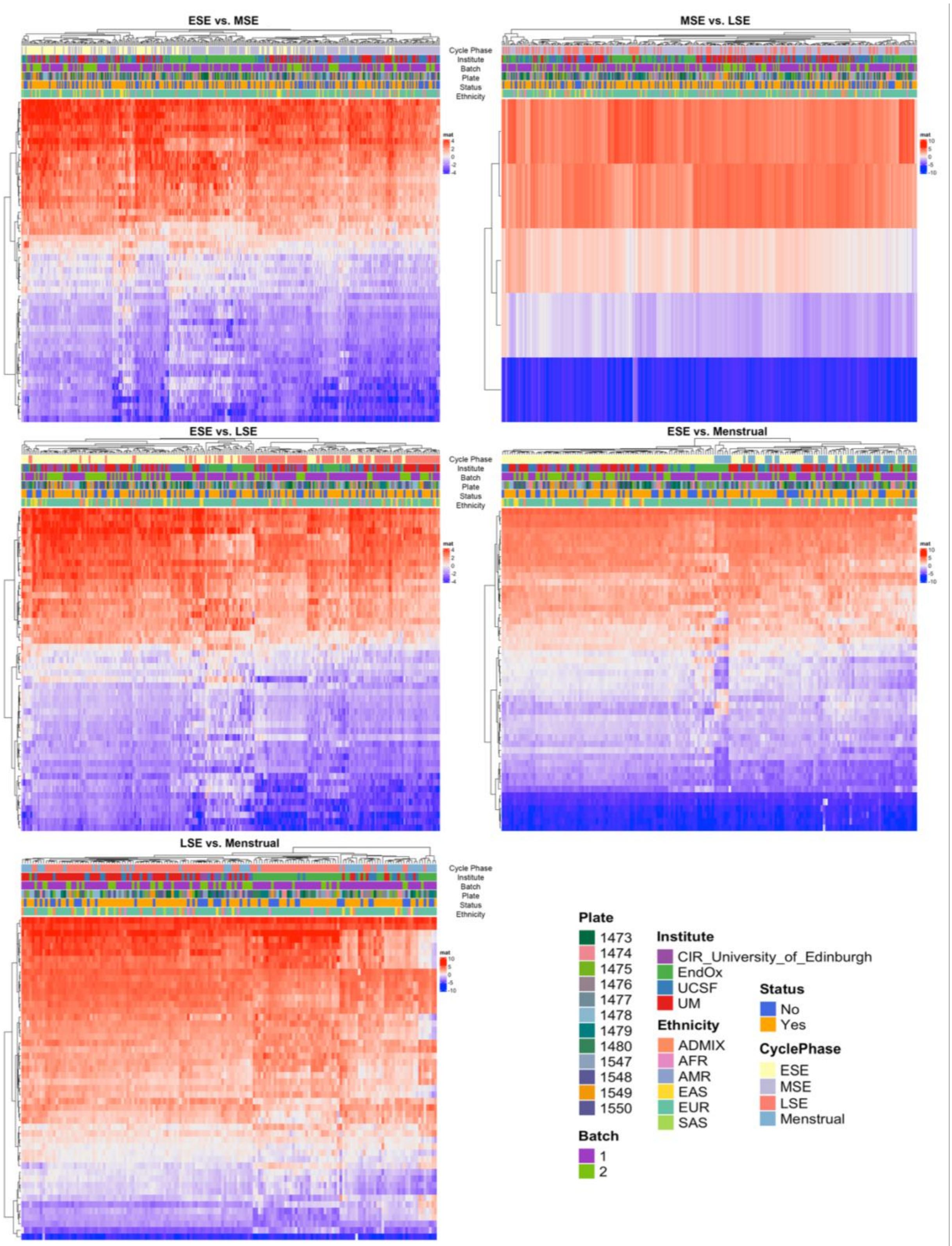


**Supplementary Figure 4. Cycle Phase Differences.** Heatmaps of the top differentially methylated DNAm sites between early secretory (ESE) and mid secretory (MSE), MSE and late secretory (LSE), ESE and LSE, ESE and menstrual phase and LSE and menstrual phase. Sample plate, batch, institute, ethnicity, endometriosis status and cycle phase are displayed as coloured bars on top of the heatmaps.

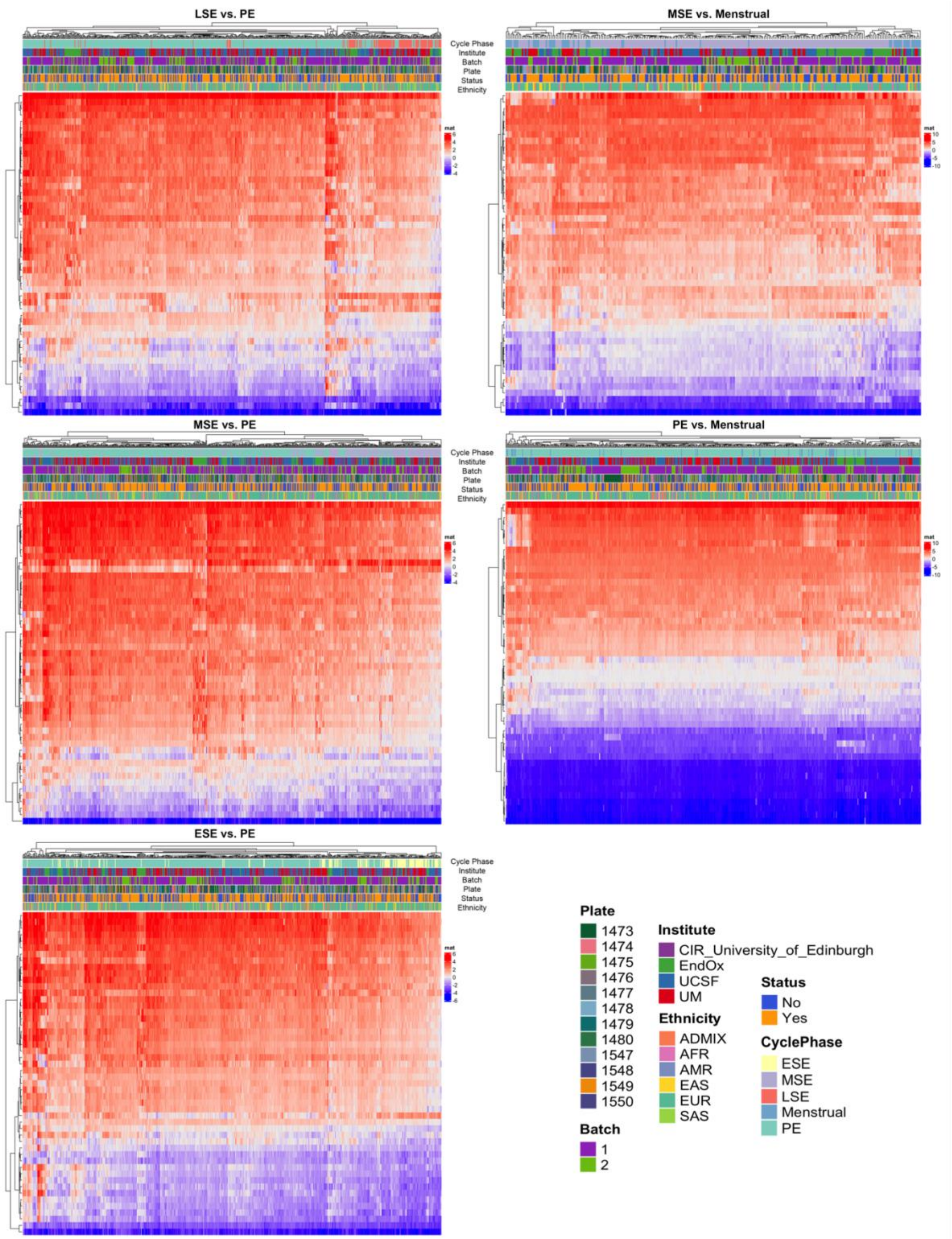


**Supplementary Figure 5. Cycle Phase Differences.** Heatmaps of the top differentially methylated DNAm sites between late secretory (LSE) and proliferative (PE), mid secretory (MSE) and menstrual phase, MSE and PE, PE and menstrual phase and early secretory (ESE) and PE. Sample plate, batch, institute, ethnicity, endometriosis status and cycle phase are displayed as coloured bars on top of the heatmaps.

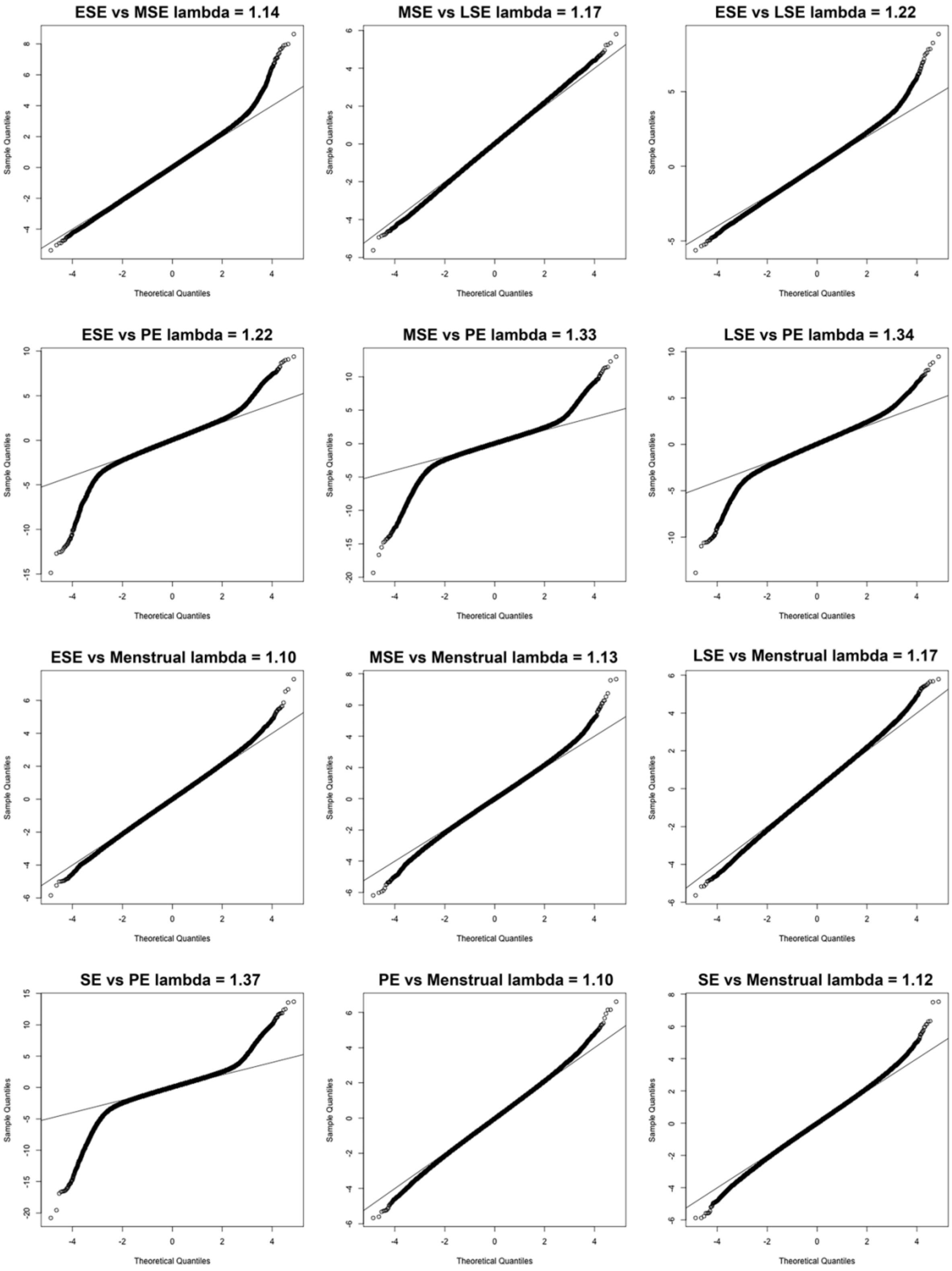


**Supplementary Figure 6. Cycle Phase Comparison QQ Plots.** QQ plots displayed the observed and theoretical quantiles for probabilities generated from 12 differential DNAm linear models comparing menstrual cycle phases (early secretory = ESE; mid secretory = MSE; late secretory (LSE); proliferative (PE) and menstrual phase).

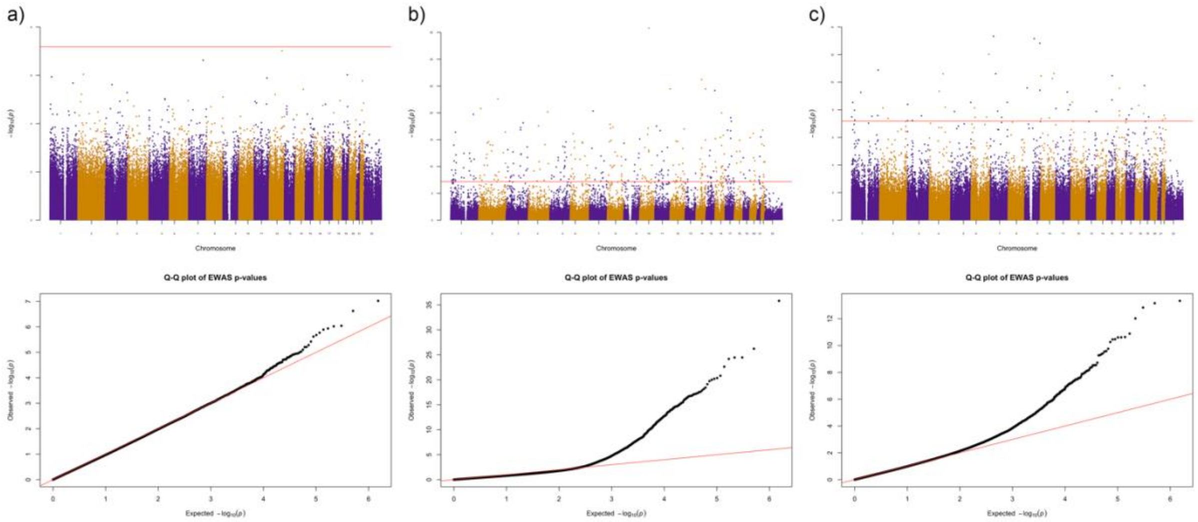


**Supplementary Figure 7. MLM-based omic association (MOA) differential DNAm.** Results of MOA for a) endometriosis cases vs controls, b) PE vs MSE and c) ESE vs LSE. The upper panels are Manhattan plots showing the association of each DNAm site as a point. Genomic location is shown on the x-axis and the -log10 p-value of the association on the y-axis. The horizontal red line represents the genome-wide significance threshold. The lower panels are QQ plots showing the expected and observed p-values for each model.

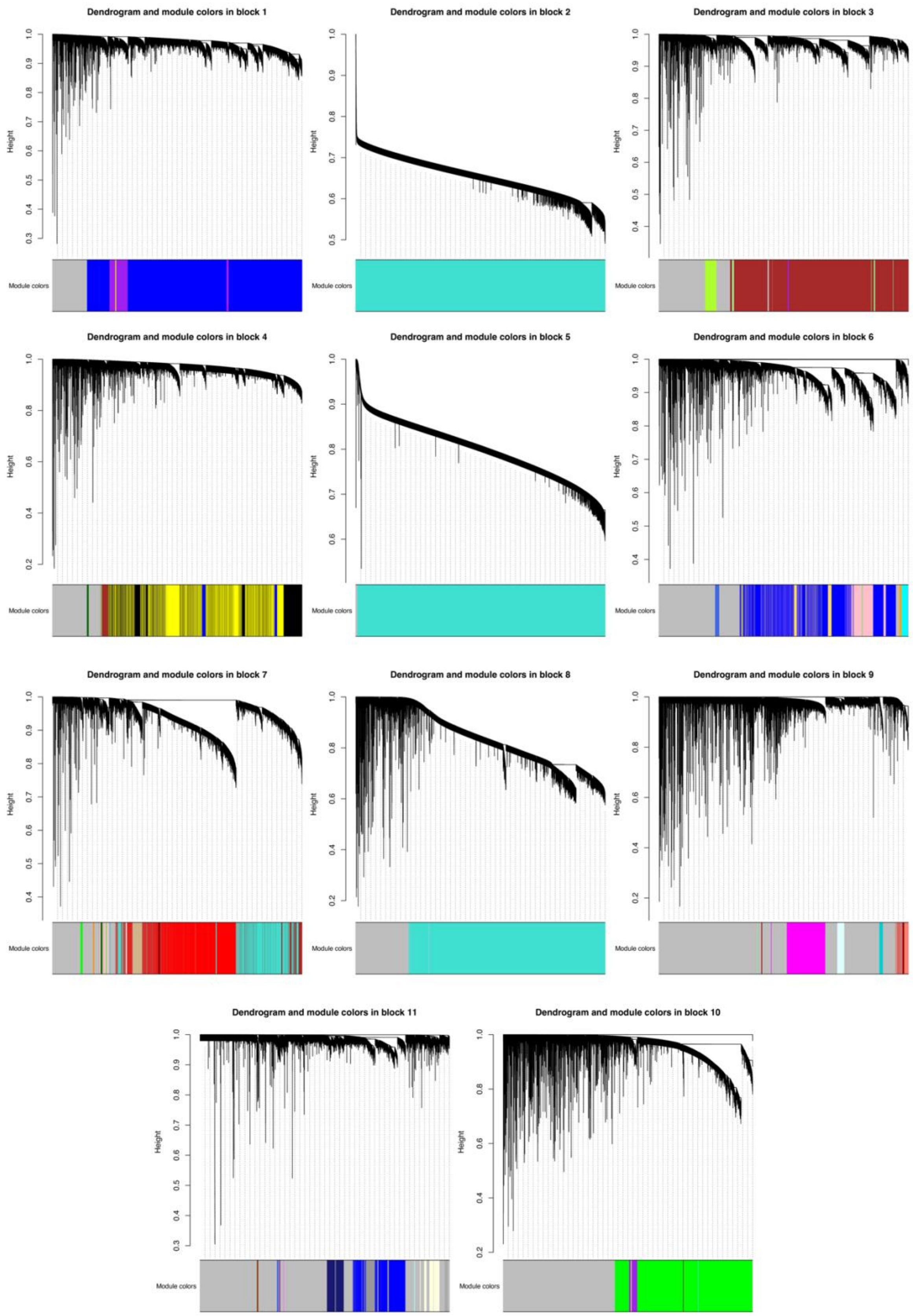


**Supplementary Figure 8. WGCNA Analysis Clustering.** Clusters (modules) of highly correlated DNAm sites generated by the Weighted correlation network analysis (WGCNA). The analysis was performed on a reduced dataset using 50% of most variable DNAm sites resulting in a dataset of 379,672 DNAm sites from the 984 samples.

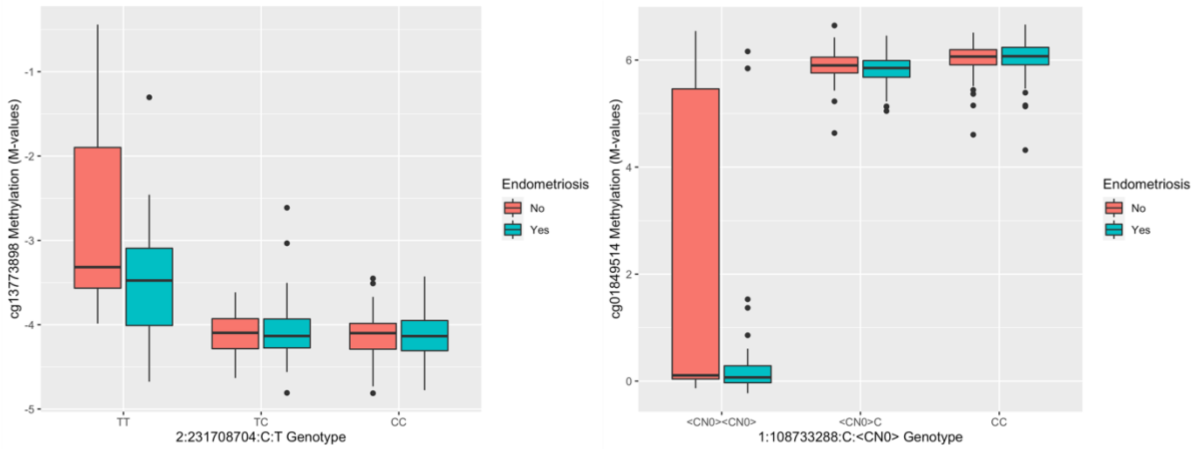


**Supplementary Figure 9. Endometriosis context specific mQTLs.** Boxplots of two mQTLs with effect sizes that are significantly different between endometriosis cases and controls.

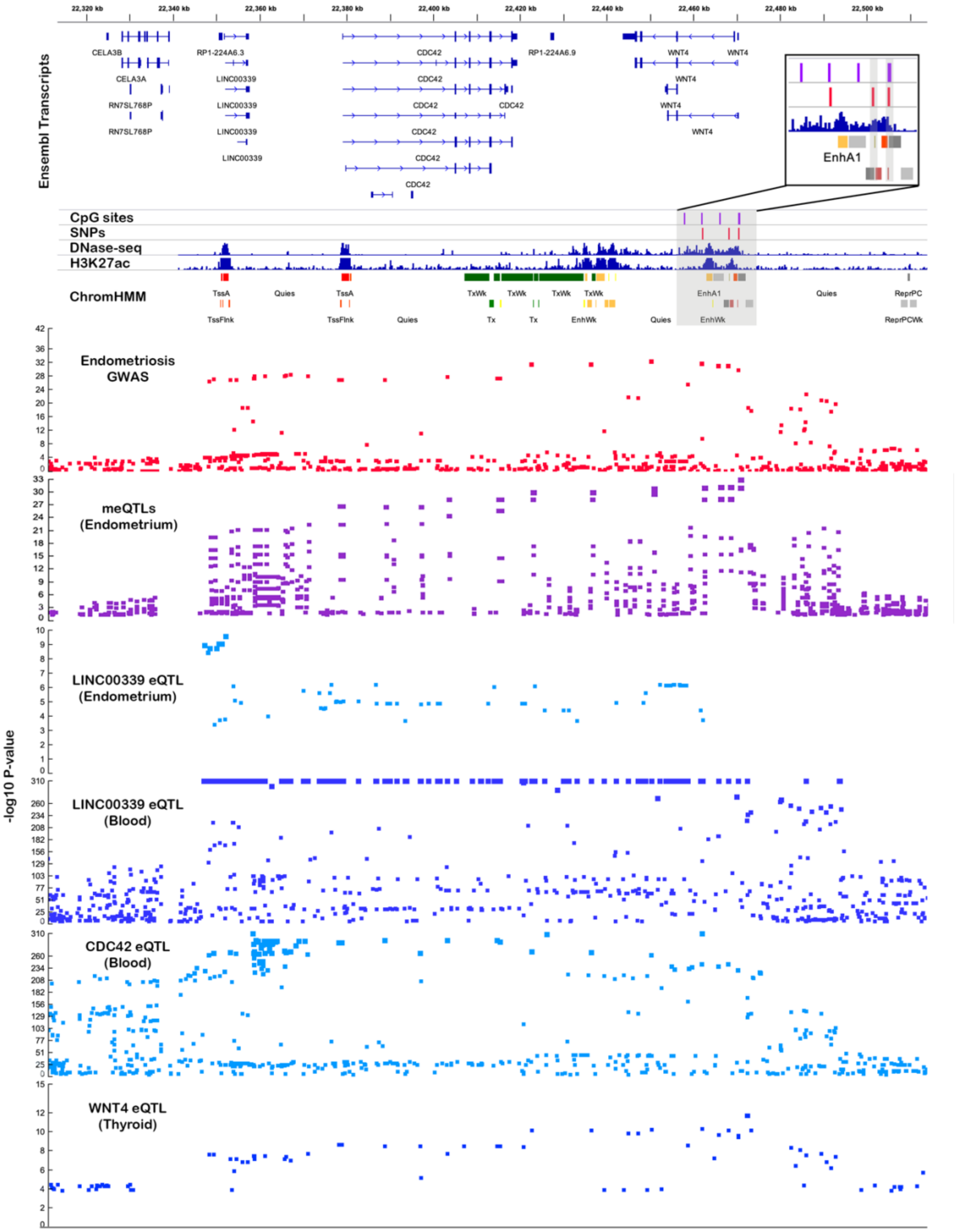


**Supplementary Figure 10. mQTLs on chromosome 1 associated with endometriosis risk.** The top panel shows ensemble transcripts present in the locus. The bottom panel consists of association plots, each point is a SNP plotted according to its genomic position on the x-axis and -log10 p-value for its association with endometriosis (red) methylation at six SMR significant CpG sites (purple) and *LINC00339*, *CDC42* and *WNT4* expression in endometrium, blood and thyroid (blue) on the y-axis. The position of the significant SMR mQTL SNPs (red)

and CpG sites (purple) is featured in the middle panel above DNase-seq peaks, H3K27ac peaks and predicted chromatin marks in uterus.

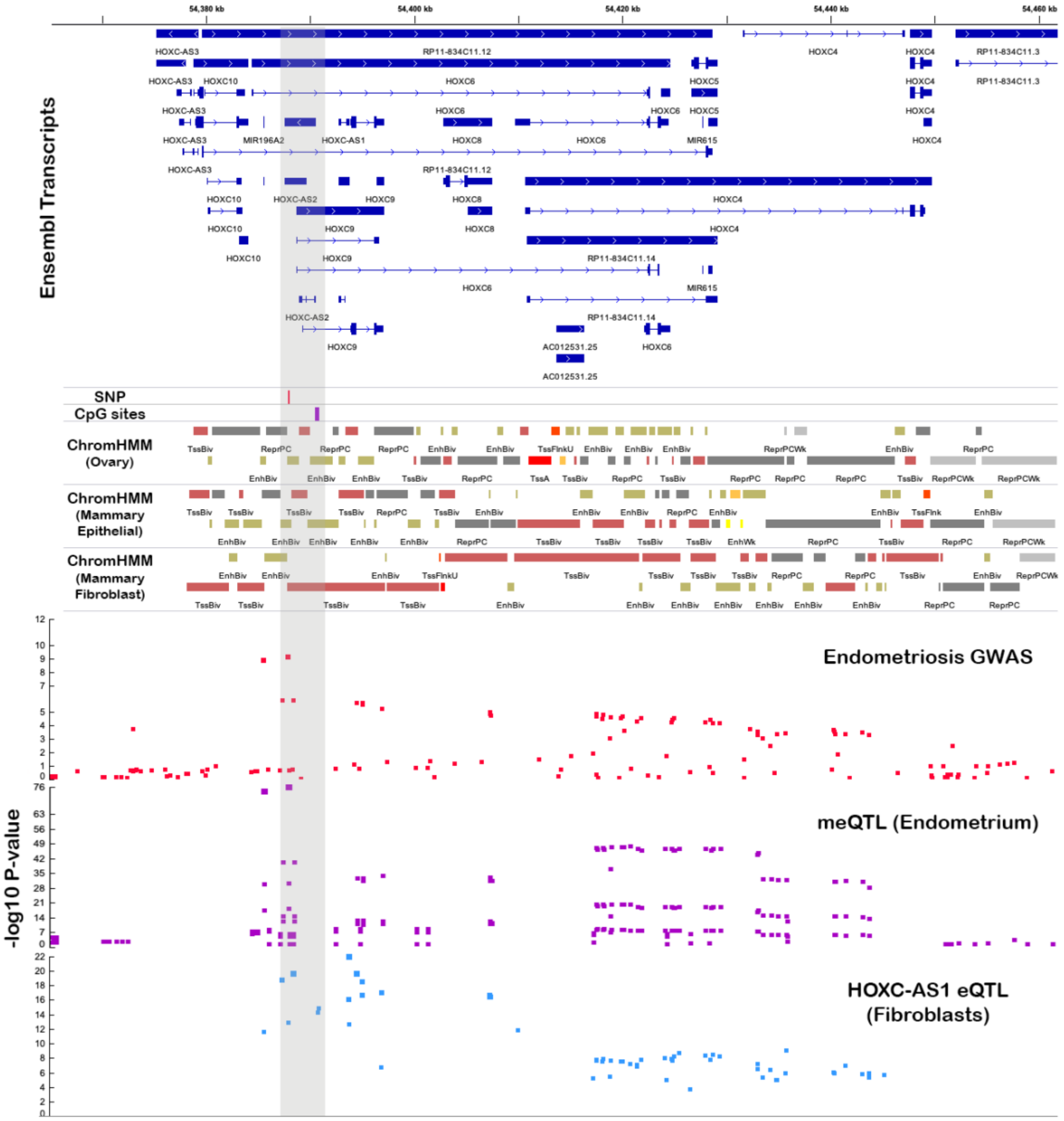


**Supplementary Figure 11. mQTLs on chromosome 12 associated with endometriosis risk.** The top panel shows ensemble transcripts present in the locus. The bottom panel consists of association plots, each point is a SNP plotted according to its genomic position on the x-axis and -log10 p-value for its association with endometriosis (red) methylation at six SMR significant CpG sites (purple) and *HOXC-AS1* expression in fibroblast cells (blue) on the y-axis. The position of the significant SMR mQTL SNPs (red) and CpG sites (purple) is featured in the middle panel above predicted chromatin marks in ovary, mammary epithelial cells and mammary fibroblast cells.

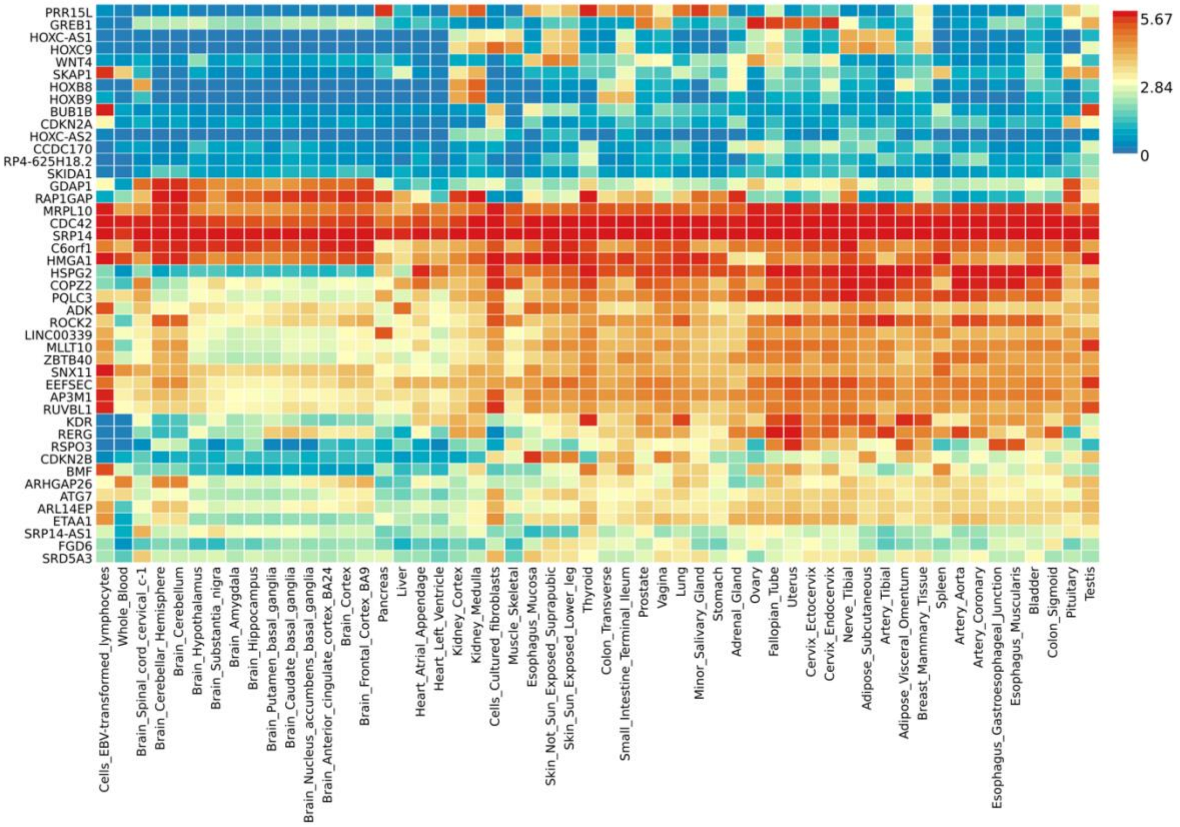


**Supplementary Figure 12. Expression of endometriosis candidate genes.** Heatmap of expression of 45 genes of interest annotated to enhancers and promoters in GTEx tissues.

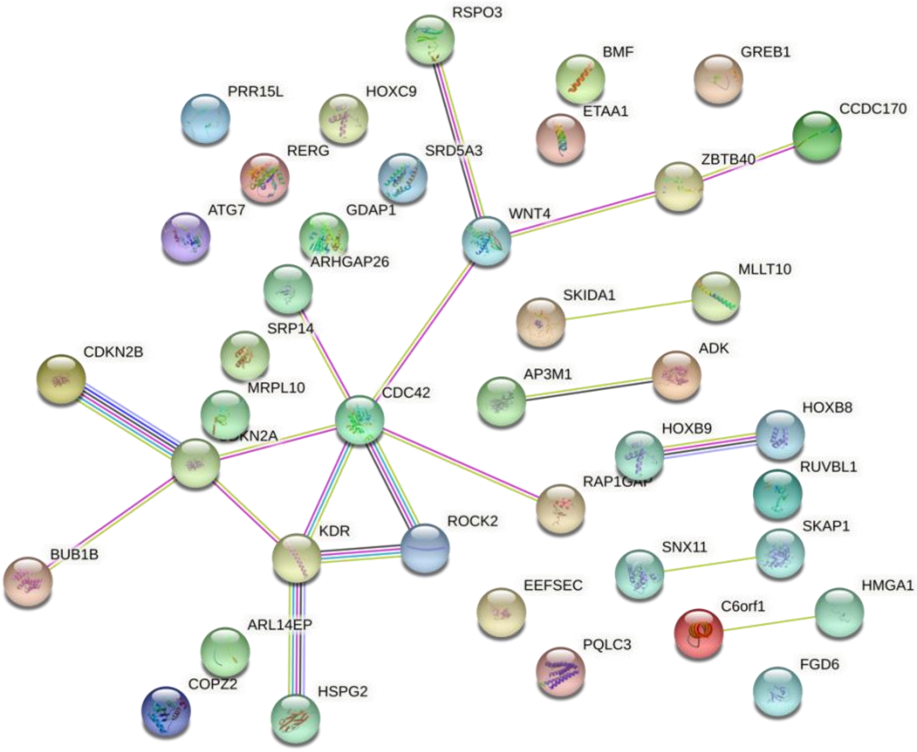


**Supplementary Figure 13. STRING protein interaction network.** Protein interactions between 45 endometriosis candidate genes of interest.

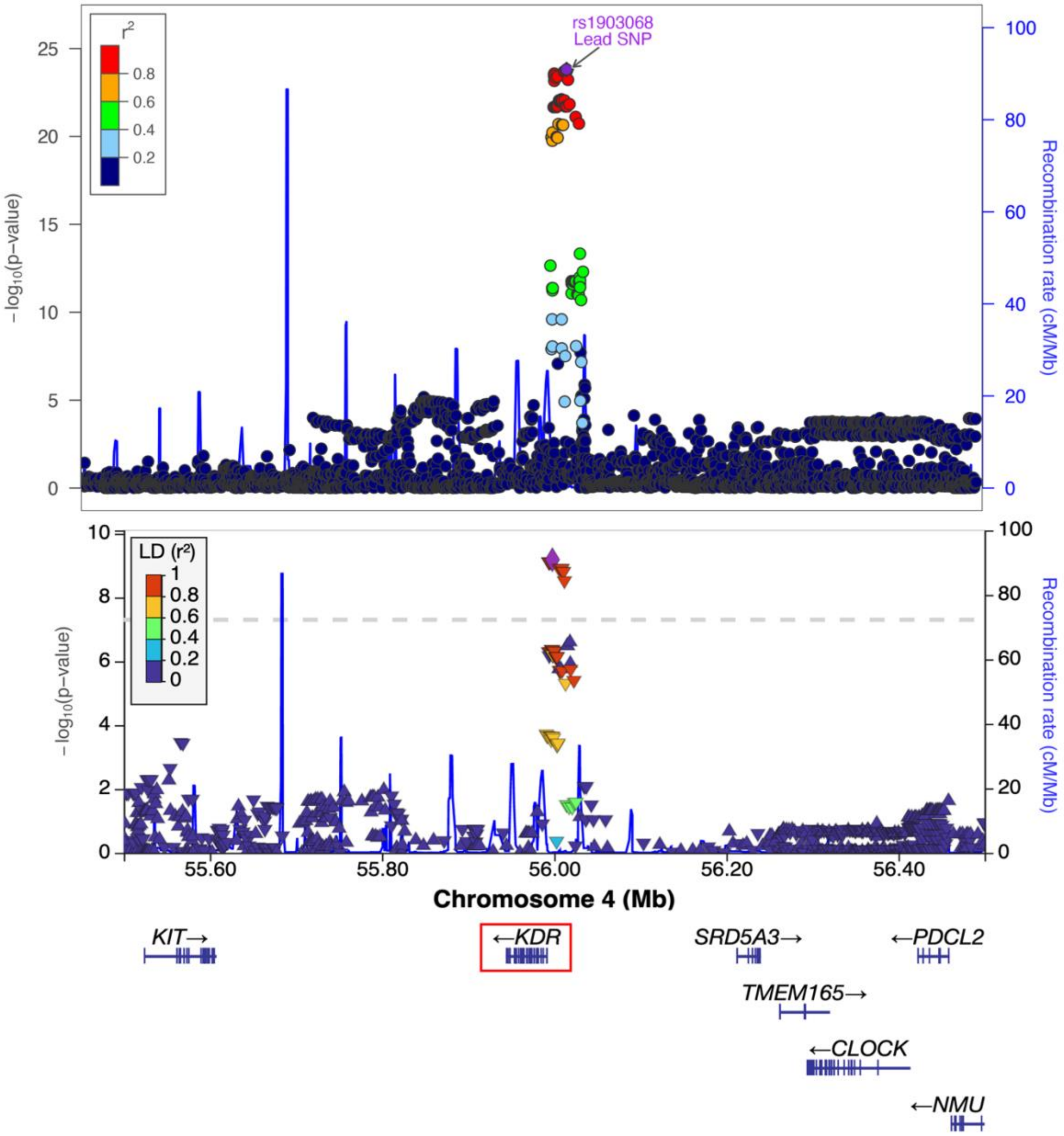


**Supplementary Figure 14. The regional association plot for 4q12.** The top panel Iillustrates the genome-association evidence of association between endometriosis and rs1903068 endometriosis lead SNP. The bottom panel illustrates the mQTL evidence from the same region highlighting the mQTL lead SNP, rs12331597 in strong LD with the endometriosis associated SNP (r2=0.97).

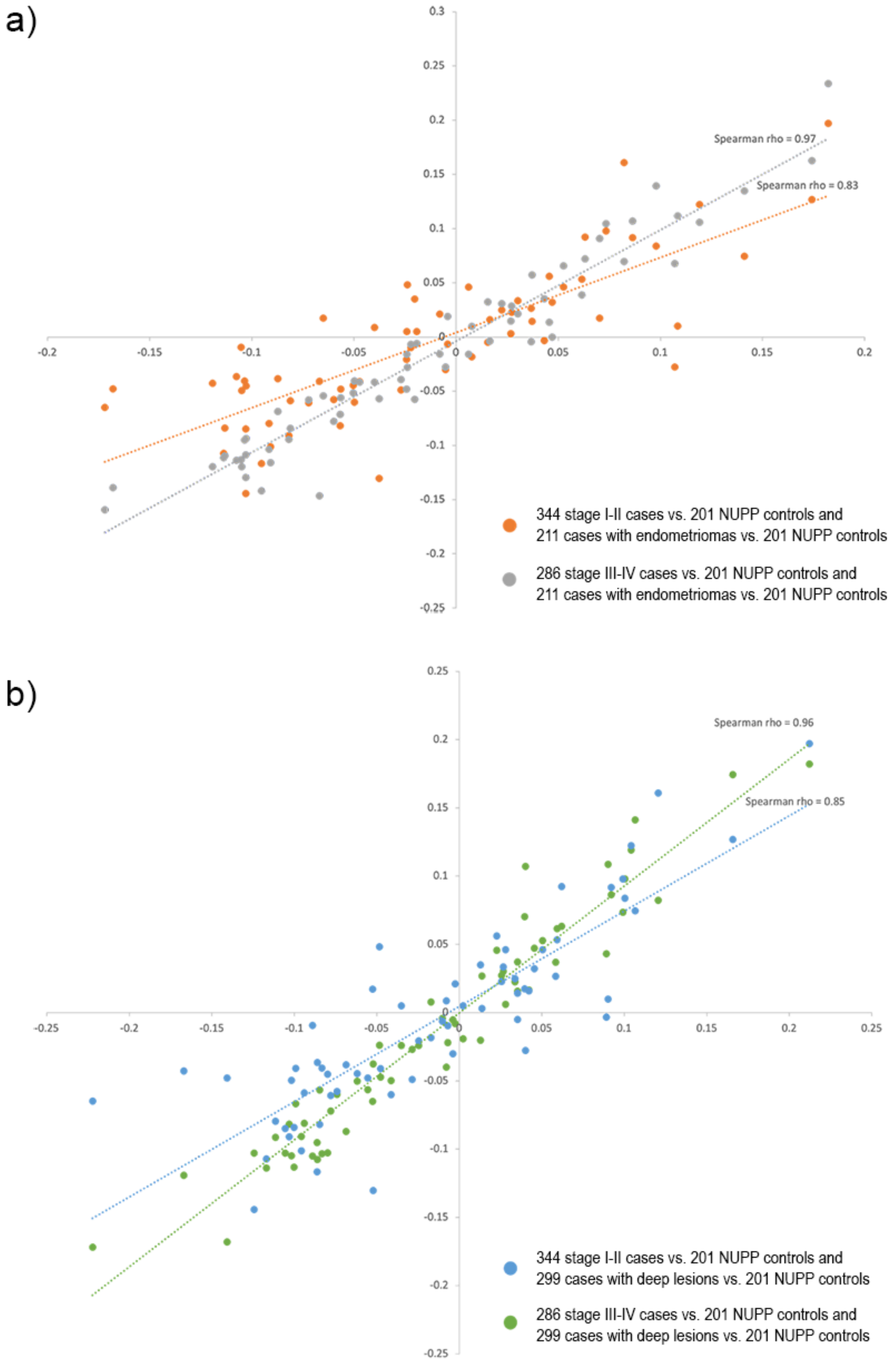


**Supplementary Figure 15.** Correlation of disease stage-based differential methylation results (the co- efficient; effect sizes), stage I/II vs. NUPP controls and stage III/IV vs. NUPP controls, with cases who have a. endometriomas and b. deep lesions.

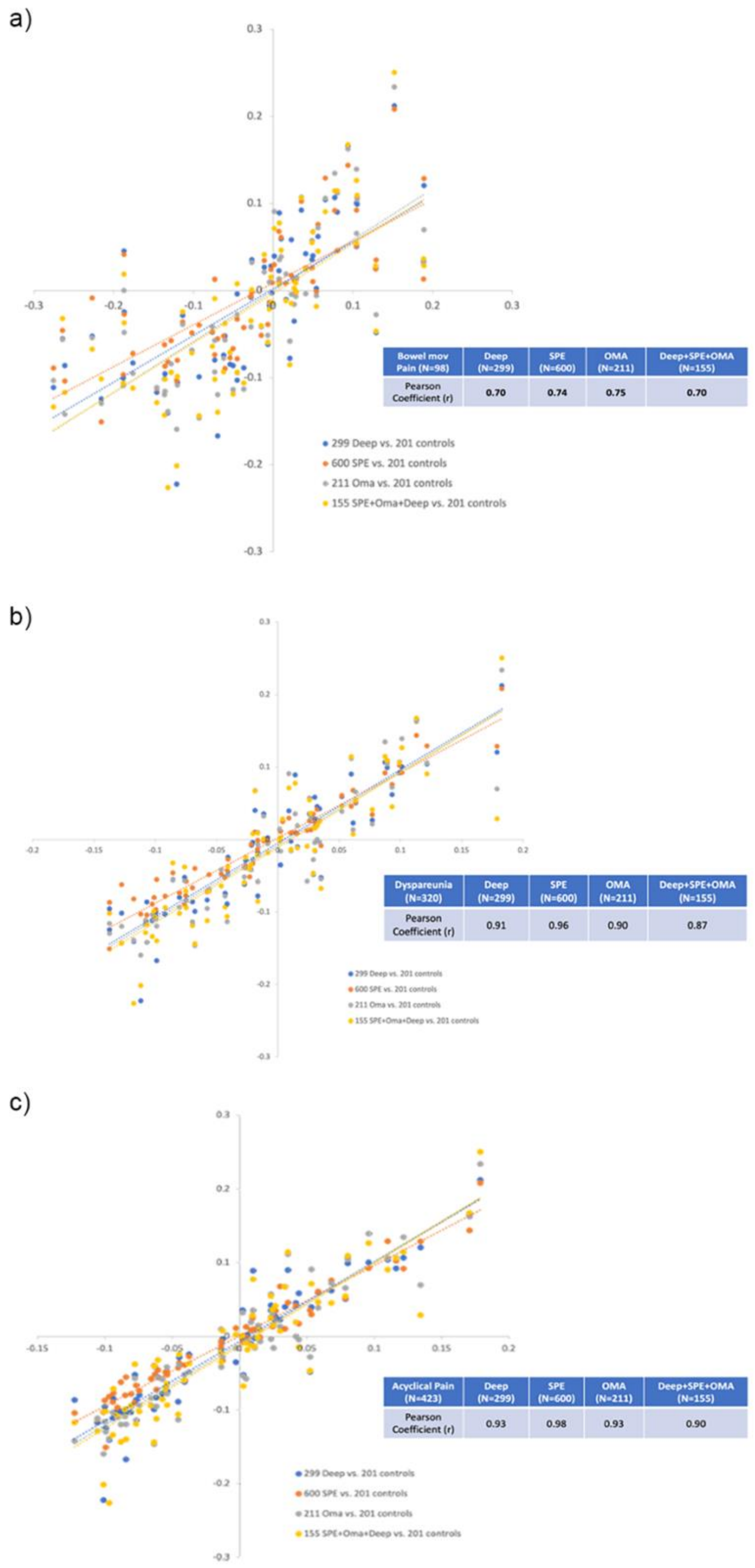


**Supplementary Figure 16.** Correlation of effect sizes of the 66 probes from the differential methylation analysis from the three main pain subtypes across the surgical subtypes including deep lesions vs. NUPP controls, superficial lesions vs, NUPP controls, endometriomas vs. NUPP controls and co-occurrence of deep, superficial and endometriomas vs. NUPP controls: a. Bowel movement pain vs. NUPP controls, b. Dyspareunia vs. NUPP controls, c. Acyclical pain vs. NUPP controls.

**Supplementary Table 4.** Genomic locations of CpG Sites significantly associated with Cycle Phase.

| Comparison | Island | N_Shelf | N_Shore | OpenSea | S_Shelf | S_Shore |
|---|---|---|---|---|---|---|
| **ESE_LSE** | 13 | 6 | 20 | 285 | 13 | 11 |
| **ESE_MSE** | 7 | 9 | 14 | 268 | 9 | 8 |
| **MSE_LSE** | 1 | 0 | 0 | 4 | 0 | 0 |
| **MSE_PE** | 178 | 262 | 389 | 6390 | 275 | 314 |
| **ESE_PE** | 63 | 102 | 149 | 2820 | 123 | 143 |
| **LSE_PE** | 155 | 105 | 193 | 2644 | 130 | 146 |
| **ESE_Menstrual** | 9 | 1 | 5 | 39 | 1 | 2 |
| **MSE_Menstrual** | 17 | 4 | 18 | 174 | 4 | 9 |
| **PE_Menstrual** | 16 | 1 | 2 | 41 | 0 | 5 |
| **LSE_Menstrual** | 3 | 1 | 7 | 57 | 1 | 4 |
| **SE_PE** | 202 | 308 | 492 | 7919 | 342 | 391 |
| **SE_Menstrual** | 16 | 4 | 15 | 117 | 2 | 9 |

**Supplementary Table 7.** Genomic Locations of CpG sites located in WGCNA modules.

| Module | Island | N_Shelf | N_Shore | OpenSea | S_Shelf | S_Shore |
|---|---|---|---|---|---|---|
| ME0 | 38 | 0 | 2 | 5 | 0 | 0 |
| ME1 | 8 | 38 | 42 | 772 | 27 | 29 |
| ME2 | 1353 | 1642 | 3809 | 39018 | 1590 | 3132 |
| ME3 | 0 | 17 | 16 | 643 | 19 | 13 |
| ME4 | 35 | 0 | 8 | 0 | 0 | 5 |
| ME5 | 575 | 1019 | 1900 | 23923 | 986 | 1664 |
| ME6 | 1025 | 728 | 1750 | 13695 | 618 | 1581 |
| ME7 | 123 | 32 | 35 | 385 | 24 | 37 |
| ME8 | 2 | 3 | 0 | 42 | 1 | 0 |
| ME9 | 0 | 6 | 6 | 130 | 4 | 3 |
| ME10 | 0 | 0 | 0 | 49 | 0 | 0 |
| ME11 | 4508 | 251 | 2381 | 3730 | 228 | 2132 |
| ME12 | 454 | 550 | 1530 | 12102 | 568 | 1310 |
| ME13 | 0 | 3 | 5 | 415 | 9 | 1 |
| ME14 | 22 | 11 | 27 | 0 | 3 | 21 |
| ME15 | 0 | 18 | 22 | 733 | 27 | 25 |
| ME16 | 4 | 44 | 135 | 1552 | 51 | 117 |
| ME17 | 0 | 23 | 19 | 816 | 17 | 13 |
| ME18 | 1311 | 368 | 1352 | 6322 | 293 | 1231 |
| ME19 | 16463 | 3132 | 8981 | 56374 | 2833 | 7704 |
| ME20 | 28 | 23 | 27 | 461 | 13 | 21 |
| ME21 | 3929 | 36 | 443 | 201 | 40 | 295 |
| ME22 | 24 | 5 | 17 | 52 | 4 | 7 |
| ME23 | 245 | 1 | 120 | 52 | 5 | 107 |
| ME24 | 113 | 19 | 98 | 401 | 22 | 77 |
| ME25 | 32 | 1 | 7 | 16 | 2 | 14 |
| ME26 | 102 | 117 | 347 | 3179 | 144 | 292 |
| ME27 | 2 | 29 | 31 | 784 | 24 | 27 |
| ME28 | 25 | 13 | 17 | 220 | 11 | 20 |
| ME29 | 0 | 0 | 1 | 48 | 0 | 1 |
| ME30 | 102 | 227 | 461 | 5502 | 233 | 404 |
| ME31 | 2502 | 4537 | 7699 | 88624 | 4379 | 6458 |
| ME32 | 987 | 18 | 91 | 164 | 16 | 81 |
| ME33 | 1 | 24 | 12 | 646 | 12 | 14 |
| ME34 | 727 | 18 | 87 | 183 | 15 | 59 |
| ME35 | 53 | 8 | 18 | 55 | 7 | 8 |

**Supplementary Note 1: Differential methylation with MLM-based omic association (MOA)**

MOA is implemented in the software package, OSCA (omic-data-based complex trait analysis)[1]. MOA fits the target probe as a fixed effect and all the other (distal) probes as random effects to account for the confounding effects, including the correlations among distal probes induced by the confounding[1]. Differences between endometriosis cases and controls were tested using the following model. M-values from the same 759,345 probes across the 984 samples were included.

Methylation ~ Endometriosis_Yes_No + Wu + e

W is an $n \times m$ matrix of standardized DNAm measures of all $m$ probes, u is an $m \times 1$ vector of the joint effects of all probes on the phenotype, and e is an $n \times 1$ vector of residuals.

MOA was also used for the largest cycle phase comparisons PE vs MSE and ESE vs LSE.

Methylation ~ Menstrual_Cycle_Phase + Wu + e

No probes were significantly differentially methylated between endometriosis cases and controls following correction for multiple testing (lambda = 0.95)(Supplementary Figure 7a). A total of 983 probes were significantly differently methylated between PE and MSE (lambda = 0.71)( Supplementary Figure 7b) 99% of which were also significant in the linear model in Limma. A total of 383 probes were significantly differently methylated between ESE and LSE (lambda = 0.96), of which only 40% were also significant in the linear model in Limma however when restricted to Bonferroni significant probes 75% were also significant in the linear model in Limma (Supplementary Figure 7c).

1. Zhang, F., Chen, W., Zhu, Z., Zhang, Q., Nabais, M.F., Qi, T., Deary, I.J., Wray, N.R., Visscher, P.M., McRae, A.F., and Yang, J. (2019). OSCA: a tool for omic-data-based complex trait analysis. Genome Biology *20,* 107. 10.1186/s13059-019-1718-z.
